# Prodomain-Growth Factor Swapping in the Structure of pro-TGF-β1

**DOI:** 10.1101/165415

**Authors:** Bo Zhao, Shutong Xu, Xianchi Dong, Chafen Lu, Timothy A. Springer

## Abstract

Transforming growth factor (TGF)-β is synthesized as a proprotein that dimerizes in the endoplasmic reticulum. After processing in the Golgi to cleave the N-terminal prodomain from the C-terminal growth factor (GF) domain in each monomer, pro-TGF-β is secreted and stored in latent complexes. It is unclear which prodomain and GF monomer are linked prior to proprotein convertase (PC) cleavage, and how much conformational change occurs following cleavage. We have determined a structure of pro-TGF-β1 with the PC cleavage site mutated, to mimic the structure of the TGF-β1 proprotein. Our structure demonstrates that the prodomain arm domain in one monomer is linked to the GF that interacts with the arm domain in the other monomer in the dimeric structure, i.e., the prodomain arm domain and GF domain in each monomer are swapped. Swapping has important implications for the mechanism of biosynthesis in the TGF-β family and is relevant to the mechanism for preferential formation of heterodimers over homodimers for some members of the TGF-β family. Our structure also provides comparisons between independent TGF-β1 crystal structures and between human and porcine pro-TGF-β1.

## Introduction

The 33 members of the transforming growth factor-β (TGF-β) family include bone morphogenetic proteins, growth and differentiation factors, activins, and inhibins. They regulate all aspects of embryogenesis, major organ development, and homeostasis [1, 2]. TGF-βs 1-3 regulate development, cell fate, wound healing, and immune responses. TGF-β protein monomers are biosynthesized with an N-terminal prodomain of ∼250 residues and a C-terminal growth factor (GF) domain of ∼110 residues [3-5]. In the endoplasmic reticulum, two monomers noncovalently associate and both the prodomain and GF domain are covalently linked into disulfide-linked dimers. At the same time, the TGF-β prodomain also associates with and becomes disulfide-linked to “milieu molecules” such as latent TGF-β binding protein (LTBP) and glycoprotein-A repetitions predominant protein (GARP) [6-9]. After transport to the Golgi, the polypeptide connection between the prodomain and GF is cleaved by a proprotein convertase. However, the GF remains strongly noncovalently bound to the prodomain, which keeps it latent during storage in extracellular milieus. Prodomain-GF interfaces are not only important for latency but also for biosynthesis. The prodomain is required for proper folding and dimerization of the GF domain [10, 11]. The C-terminal GF domain folds either concomitantly with, or subsequently to, the N-terminal prodomain [10, 12, 13].

The structure of the pro-TGF-β1 dimer reveals that the prodomain dimer surrounds the GF dimer [14]. Two arm domain monomers with a jelly roll, β-sandwich fold dimerize and are disulfide linked at a bowtie knot and surround the GF on one side. On the other side, the N and C-termini of the prodomain form a straitjacket that more loosely surround the GF. Each straitjacket monomer contains an α1-helix that intercalates between the two GF monomers, a latency lasso that wraps around the distal ends of each GF monomer, and an α2-helix that snuggles against the GF-arm domain interface. TGF-β cannot bind its receptors and signal until it is released from the prodomain, i.e. activated [15-17]. TGF-βs 1 and 3 are activated by integrins that bind to a RGDLXXI/L motif in the prodomain [18]. The motif locates to a long loop in the arm domain called the bowtie tail, because it follows the two bowtie cysteines that disulfide link the two prodomain monomers. A structure of an α_V_β_6_ integrin head bound to one monomer of a TGF-β dimer showed that integrin binding stabilizes an alternative conformation of the bowtie tail [19]. Activation by integrin α_V_β_6_ requires force application by the actin cytoskeleton, which is resisted by the milieu molecule, resulting in distortion of the prodomain and release of the GF [9, 15, 16, 19].

The dimeric pro-TGF-β1 crystal structure suggested that there might be a swap between the prodomain and GF domain [14]. In the structure, each prodomain interacts much more with one monomer than the other. The solvent accessible surface areas buried between one prodomain and two different GF monomers are 900 Å^2^ and 370 Å^2^, respectively. The larger of the two prodomain-GF interfaces is larger than the interface between the two prodomains (600 Å^2^) or between the two GF domains (330 Å^2^). The structure that suggested swapping was of mature pro-TGF-β1, which was cleaved between each prodomain and GF monomer. The C-terminus of one prodomain was closer to the N-terminus of one GF monomer than the other, and if these were linked in the original monomer, this monomer would correspond to the prodomain and GF monomers with the smallest amount of noncovalent association in the final structure [14]. In other words, there would have been a swap. However, it was unknown how closely the structure of PC-cleaved, pro-TGF-β1 resembled that of the pro-protein prior to PC cleavage. Cleavage might have allowed substantial structural rearrangements. Furthermore, the longer connection could not be ruled out structurally.

Here, we have determined a structure of pro-TGF-β1 with the PC cleavage site mutated, to mimic the structure of the TGF-β1 pro-protein prior to PC cleavage. Our structure supports prodomain-GF swapping. Swapping also has important implications because it can provide a mechanism for preferential formation of heterodimers over homodimers when a cell synthesizes monomers for two different TGF-β family members. Such heterodimers can display unique biological activities as in the case of BMP-2/7 heterodimers [20], and can alter activity as in the case of inhibin heterodimers compared to activin homodimers [12].

We also describe differences among pro-TGF-β1 crystal structures that appear unrelated to prodomain cleavage and relate to the issue of which regions in pro-TGF-β1 are flexible. Flexibility is relevant to TGF-β activation because rigidity is important for force transmission, and flexibility is important for release of the GF from the prodomain and for the ability of pro-TGF-β1 to covalently and noncovalently associate with structurally distinct milieu molecules [6-9].

## Results

### Crystal structure of pro-TGF-β1 prior to furin cleavage

We introduced an R249A mutation into the RHRR^249^ PC cleavage site between the prodomain and GF domain of human pro-TGF-β1. SDS-PAGE showed that the R249A mutant is >90% uncleaved whereas WT protein is almost completely cleaved (Fig.1A). Reducing SDS-PAGE of R249A mutant crystals showed a band corresponding to uncleaved pro-TGF-β1 monomer (Fig. 1B), while porcine pro-TGF-β1 crystals show predominantly the cleaved prodomain [14].

**Figure 1:**
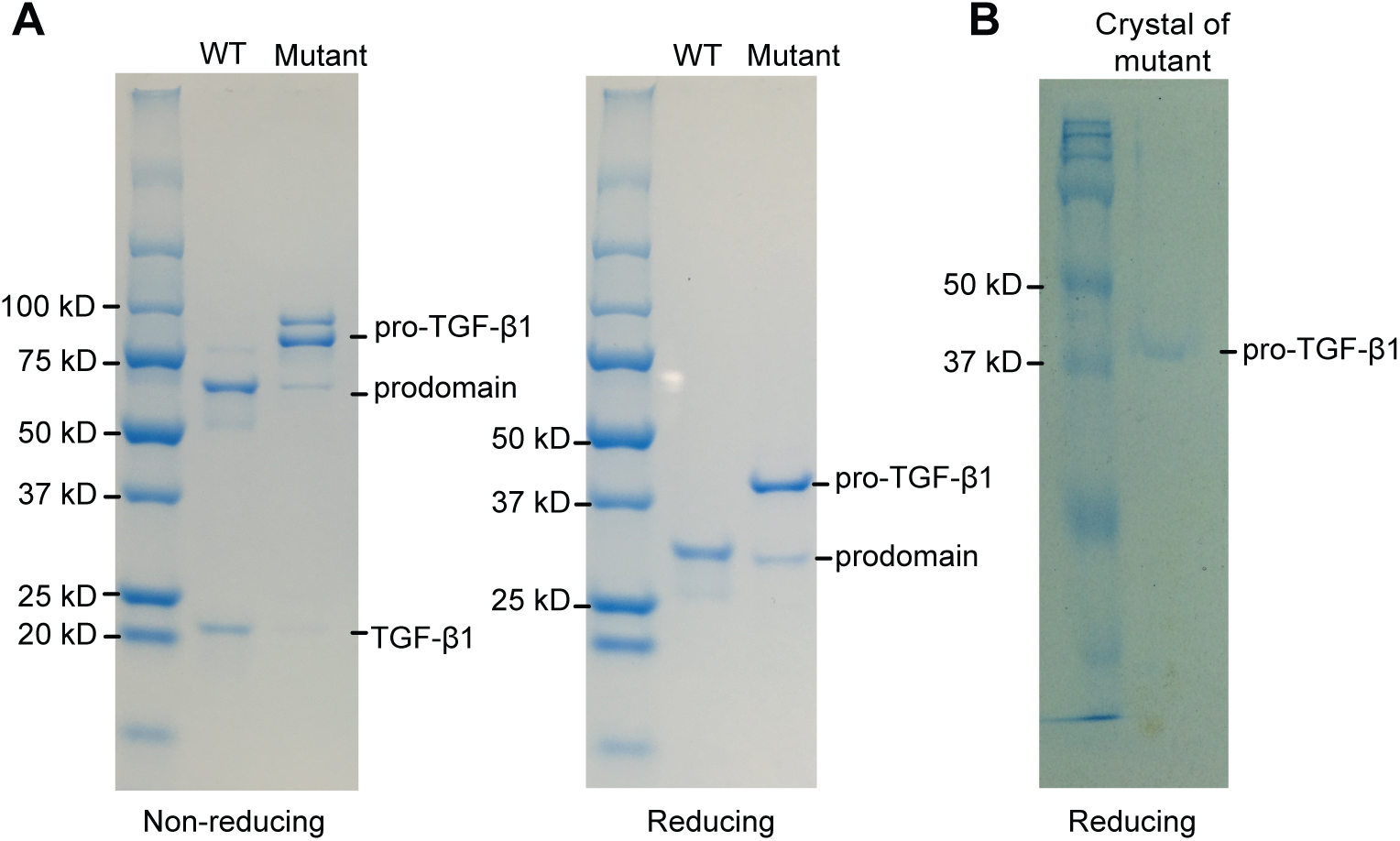
SDS-PAGE of purified pro-TGF-β 1 and crystal. **A**. Non-reducing (left) and reducing (right) SDS 4-20% PAGE of purified WT and R249A mutant pro-TGF-β1. **B.** Reducing SDS-PAGE (10%) of a crystal of pro-TGF-β1 R249A mutant. Gels were stained with Coomassie Brilliant Blue.

We determined a crystal structure to a resolution of 2.9 Å of human R249A mutant pro-TGF-β1 (Table 1, Fig. 2A). The structure contains one monomer in the asymmetric unit; the other monomer of the dimer shown in Figures is a symmetry mate. We also re-processed to higher resolution and re-refined the previously described, cleaved porcine pro-TGF-β1 structure (Fig. 2B). Despite the increase in resolution to 2.9 Å, the Rfree is lower than for the previous model (Table 1). The porcine, PC-cleaved structure has four monomers in the asymmetric unit but nonetheless has pseudosymmetry that resembles the symmetry of the human R249A mutant and packs similarly in the crystal lattice.

**Table 1:** Statistics of X-ray diffraction and structure refinement

| | pro-TGF-β1 WT (5VQF) | pro-TGF-β1 R249A mutant (5VQP) |
| --- | --- | --- |
| **Data collection statistics** |  |  |
| Space group | P 2<sub>1</sub> | P 6 2 2 |
| $\alpha, \beta, \gamma, ^\circ$ | 90, 96.7, 90 | 90, 90, 120 |
| Unit cell (a, b, c), Å | 54.7, 126.9, 137.9 | 104.4, 104.4, 141.9 |
| Resolution range (Å) | 50.0-2.9 (3.0-2.9)<sup>a</sup> | 50.0-2.9 (3.0-2.9)<sup>a</sup> |
| Completeness (%) | 96.1 (97.8) | 99.7 (99.9) |
| Number unique reflections | 39,918 (4,031) | 10,672 (1,024) |
| Redundancy | 4.3 (4.3) | 10.5 (10.9) |
| Rmerge<sup>b</sup> (%) | 5.2 (274) | 17.1 (601) |
| I/$\sigma$ | 15.3 (0.6) | 12.3 (0.5) |
| CC<sub>1/2</sub> (%)<sup>c</sup> | 99.9 (32.6) | 99.9 (15.1) |
| Wavelength (Å) | 1.0332 | 1.0332 |
| **Refinement statistics** |  |  |
| Rwork<sup>c</sup> (%) | 23.9 (45.2) | 25.2 (41.3) |
| Rfree<sup>c</sup> (%) | 28.0 (52.0) | 29.3 (42.3) |
| Bond RMSD (Å) | 0.005 | 0.004 |
| Angle RMSD (°) | 0.85 | 0.64 |
| Ramachandran plot<sup>d</sup> |  |  |
| (Favored/allowed/outlier) | 96.7/3.3/0 | 95.5/4.5/0 |
| MolProbity percentile<sup>e</sup> |  |  |
| (Clashscore/Geometry) | 98/100 | 96/99 |
| No. atoms |  |  |
| protein | 10658 | 2576 |
| carbohydrates | 95 | 39 |
| Number of cis-Proline | 4 | 1 |
| B factors |  |  |
| Protein | 188.4 | 127.0 |
| carbohydrates | 266.4 | 221.8 |

a. The numbers in parentheses refer to the highest resolution shell.
b. $R_{\text{merge}} = \sum_h \sum_i |I_i(h) - \langle I(h) \rangle| / \sum_h \sum_i I_i(h)$ , where $I_i(h)$ and $\langle I(h) \rangle$ are the $i^{\text{th}}$ and mean measurement of the intensity of reflection $h$ .
c. Pearson's correlation coefficient between average intensities of random half-datasets for unique reflection [34].
d. $R_{\text{factor}} = \sum_h ||F_{\text{obs}}(h)| - |F_{\text{calc}}(h)|| / \sum_h |F_{\text{obs}}(h)|$ , where $F_{\text{obs}}(h)$ and $F_{\text{calc}}(h)$ are the observed and calculated structure factors, respectively. No $I/\sigma(I)$ cutoff was applied.
e. Calculated with MolProbity [22].

**Figure 2:**
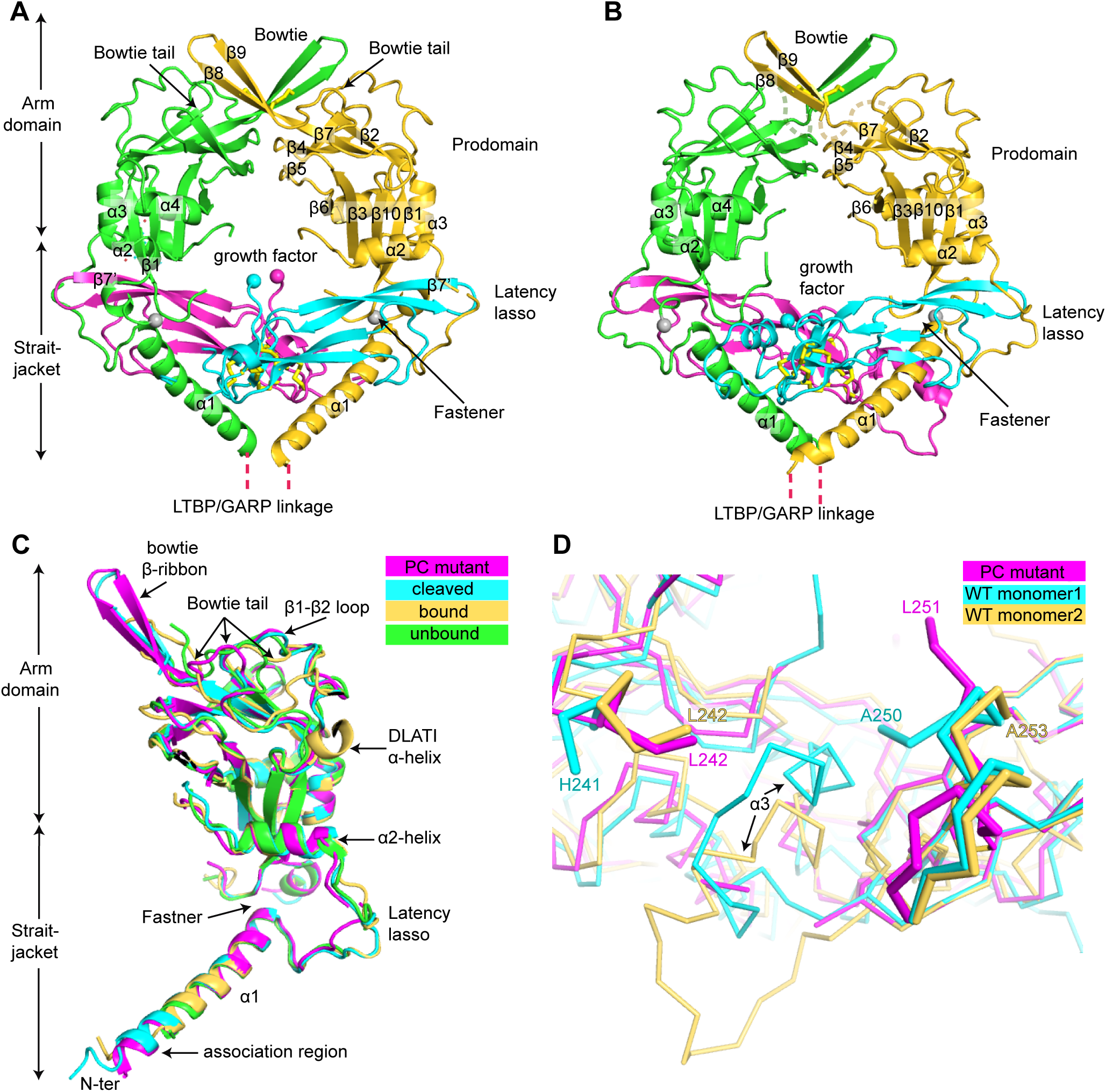
Crystal structures and comparisons. **A-B.** Crystal structure of pro-TGF-β1 R249A mutant (A) and re-refined crystal structure of WT cleaved porcine pro-TGF-β1 (B). Ribbon cartoon is colored differently for each domain. Disulfide bonds (yellow) are shown in stick. C-termini of the prodomain and the N-termini of the GF are shown as spheres. **C.** Comparison of 4 different pro-TGF-β1 prodomain monomers: the human R249A mutant, porcine WT, and the integrin-bound and unbound monomers in the 1:2 integrin: pro-TGF-β1 complex structure [19]. **D.** Comparison of the GF N-termini of different crystal structures. R249A PC mutant, and WT porcine cleaved monomers 1 (chain D) and 2 (chain C).

We first describe overall structure before focusing on structural alterations around the PC cleavage site. Comparisons give insights into regions of pro-TGF-β1 that are flexible, and hence may be affected by lattice contacts in crystals, as opposed to regions of pro-TGF-β1 that are structurally conserved. Flexibility is relevant to pro-TGF-β1 biology including integrin-dependent reshaping of the bowtie tail, reshaping of the straitjacket region by applied force to release the GF in TGF-β1 activation, force transmission from the integrin binding site through the arm domain to the straitjacket, and the ability of the N-terminal portion of the prodomain to noncovalently associate with and disulfide link to structurally distinct milieu molecules [6-9]. We compare four pro-TGF-β1 monomers, one from each of the structures refined here and two corresponding to integrin-bound and free monomers from a structure of dimeric pro-TGF-β1 with one of its monomers complexed with α_V_β_6_ [19](Fig. 2C).

In the prodomain arm domain, β-strands β1 to β7 of the jelly roll β-sandwich fold are structurally invariant among the four representative monomers, as are major α-helices α2 to α4 of the arm domain (Fig. 2A,B), as shown in the overlay of the four representative structures (Fig. 2C). β-sheet domains are relatively resistant to force. The relatively rigid arm domain β-sandwich fold thus enables integrins bound to the RGDLATI motif to transmit force through the arm domain to the straitjacket elements that surround the TGF-β1 GF [19].

In contrast, the remarkably long, 30-residue bowtie tail that contains the integrin-binding RGDLATI motif is markedly variable. When integrin is bound, the DLATI portion of this motif forms an α-helix, and protrudes from one end of the arm domain distal from the GF domain (Fig. 2C). Bowtie tail residues that bind in a hydrophobic groove in the arm domain shift by 7-11 residues in position in the sequence but assume a similar backbone conformation in the groove [19]. In the human PC-mutant monomer described here, all residues in the bowtie tail can be traced, whereas 6 to 9 residues are disordered in previous structures. However, density is poor in some regions of the PC-mutant bowtie tail, consistent with large variation between the four structures, including the previous porcine structure which crystallized in a similar lattice environment as the human R249A mutant described here. The 2-stranded bowtie β-ribbon in these two structures (Fig. 2A-C) is a consequence of formation of a 4-stranded super β-sheet with a 2-stranded bowtie β-ribbon from a neighboring molecule related by symmetry or pseudo-symmetry. In the absence of such a lattice contact, no β-ribbon is present and 9 residues are disordered in the integrin-unbound monomer from the α_V_β_6_ complex [19]. Notably, the β1-β2 loop that neighbors the bowtie tail is also highly structurally variable (Fig. 2C). This loop corresponds to a long meander between arm domain β-strands 1 and 2. Backbone movement in the meander should facilitate reshaping of the neighboring bowtie tail.

In the straitjacket, the C-terminal portion of the prodomain α1-helix, which inserts between the two GF domain monomers, is highly conserved in position. The C-terminal end of the α1-helix interacts with residues 74-76 to form a fastener that surrounds one GF monomer; each portion of the fastener is also highly conserved in position (Fig. 2C). The α2-helix, which interacts with both the GF and the arm domain, is also highly conserved structurally among the four representative pro-TGF-β1 monomers. This conservation in position of the two fastener elements and the α2-helix correlates with the observation that they are the most force-resistant straitjacket elements when integrin and actin cytoskeleton-dependent pulling on pro-TGF-β1 is resisted by the α1-helix residues that link to milieu molecules; i.e. fastener and α2-helix disruption correlates with peaks in force during pulling [19].

In contrast, the latency lasso and N-terminus of the prodomain are highly variable among the four representative straitjacket structures. In the human R249A mutant pro-TGF-β1 structure, the latency lasso is displaced up to 7 Å by a lattice contact relative to other structures and two residues are disordered. Almost the entire latency lasso, residues 33-45, differs by 2 Å or more among the four structures, showing that the lasso only loosely wraps around the GF. At the N-terminus, residues 1-2 are disordered in the human R249A mutant. The following α1-helix is stabilized as α-helical by contacts with the α1-helix of a symmetry-related molecule in the crystal lattice, as also occurred in in the previous porcine, PC-cleaved pro-TGF-β1 structure [14]. In the absence of such lattice contacts in the integrin complex with pro-TGF-β1, residues 1-9 are disordered or differ in conformation [19]. Prodomain residues 1-9 have no contact with other portions of pro-TGF-β1 and correspond to an “association region” that contains Cys-4 that disulfide links to LTBP or GARP. Because these milieu molecules have no structural similarity, the association region must be able to adopt different conformations to noncovalently associate with them. Furthermore, one pro-TGF-β1 dimer associates with a single milieu molecule in asymmetric 2:1 complexes. Thus the association regions of the two monomers bound to a milieu molecule must adopt different conformations to bind to distinct portions of a milieu molecule.

### The structure around the PC cleavage site

The conformation of the growth factor domain N-terminus in the R249A mutant pro-TGF-β1 monomer differs markedly from all four crystallographically distinct cleaved pro-TGF-β1 monomers (Fig. 2A, B, D). In the uncleaved, R249A mutant we can trace the electron density for the polypeptide chain up to prodomain residue 242. After 7 prodomain residues and the N-terminal GF residue missing in density, the trace of the GF begins with its second residue, residue 251. Residue 251 is present in the solvent-filled cavity between the arm domains and the GF in the center of the ring-shaped pro-TGF-β1 dimer (Fig. 2A, D). Residues 251-253 lack any stabilizing van der Waals or hydrogen bond interactions with neighboring residues. The ordering of residues 251-253 in the absence of any interactions with visualized residues suggests that their position is stabilized by the linkage of residue 251 to disordered residues 243-250, which connect to ordered prodomain residue 242.

In the mature, cleaved GF, the N-terminus has a distinct conformation (Fig. 2D). Residues 252-258 in the mature GF are α-helical, in contrast to the much more extended conformation in the uncleaved GF. Although the conformations differ among the N-termini of each of the four crystallographically distinct GF monomers in cleaved, WT pro-TGF-β1, and only two of four examples show ordering of N-terminal residue 250, their structural environments are overall similar in having close interactions with other GF residues and none extend into the solvent filled channel in the middle of the pro-TGF-β1 ring (Fig. 2D). The N-terminus of the mature GF extends towards the α3-helix of the GF, which is highly variable in position among the four examples of the mature GF in TGF-β1 pro-complexes and is disordered in the uncleaved GF in R249A mutant pro-TGF-β1. In concert with movement of the α3-helix, the N-terminus of the mature GF moves so that it remains in van der Waals contact with the GF. As a consequence of the extension of residues 251-254 in uncleaved pro-TGF-β1 toward the solvent channel in the ring instead of toward the α3-helix, residue 251 adopts positions that differ by 8-9 Å in uncleaved compared to cleaved pro-TGF-β1 (Fig. 2D).

The overall structures of WT and R249A mutant pro-TGF-β1 show that there are no large-scale conformational changes in the prodomain or GF or their orientation with respect to one another that relate to separation of the prodomain from the GF by PC cleavage. The most significant difference localizes to the N-terminus of the GF domain, which as described above, is altered in orientation in absence of cleavage. Eight residues (243-250) linking the prodomain to the growth factor domain are missing in density, consistent with the need of flexibility to fit into the catalytic site of the PC protease.

### Evidence for arm domain and growth factor swapping

We used modeling and mutagenesis to establish which prodomain-GF connection was physiologic. The most C-terminal residue with electron density in one prodomain (labeled C-ter 1 in Fig. 3A) must link through disordered residues either to the N-terminus of one GF monomer (N-ter 1, a distance of 18 Å) or to the N-terminus of the other monomer (N-ter 2, a distance of 32 Å). Whereas a direct connection between C-ter 1 and N-ter 1 can be built over the 18 Å distance, the connection to N-ter 2 of 32 Å cannot be built straight, because fastener residues 72-74 and the GF dimer are in the way, and the polypeptide must curve around them as it goes through the central cavity of pro-TGF-β to connect. To test feasibility of the alternative arm domain-GF connections, we built the residues missing in density using Modeller [21]. As reported by MolProbity [22], the 32 Å distance could not be spanned without introducing substantial clashes, bad geometry and bond length outliers, i.e., chain breaks. The 32 Å connection showed 100% bond length outliers over residues 242-251 for 20 of 20 models. As a further test, validation for deposition to the protein databank reported that the 32 Å connection was too long to be spanned for the number of missing residues. Moreover, lack of flexibility in the 32 Å connection would prohibit the remodeling required to fit into the PC protease active site cleft, and would position the cleavage site in the center of the cavity of the pro-TGF-β ring, where it would be inaccessible to the large, 90 kDa PC protease. In contrast, the 18 Å distance could easily be spanned by multiple conformations of the prodomain-GF linker, including those with the RHRR^249^ PC recognition motif well exposed to solvent (Fig. 3B). Representatives of the 20 connections built by Modeller between C-ter 1 and N-ter 1 are shown in different colors in Fig. 3B, with the four residues in the PC recognition motif shown as small Cα-atom spheres.

**Figure 3:**
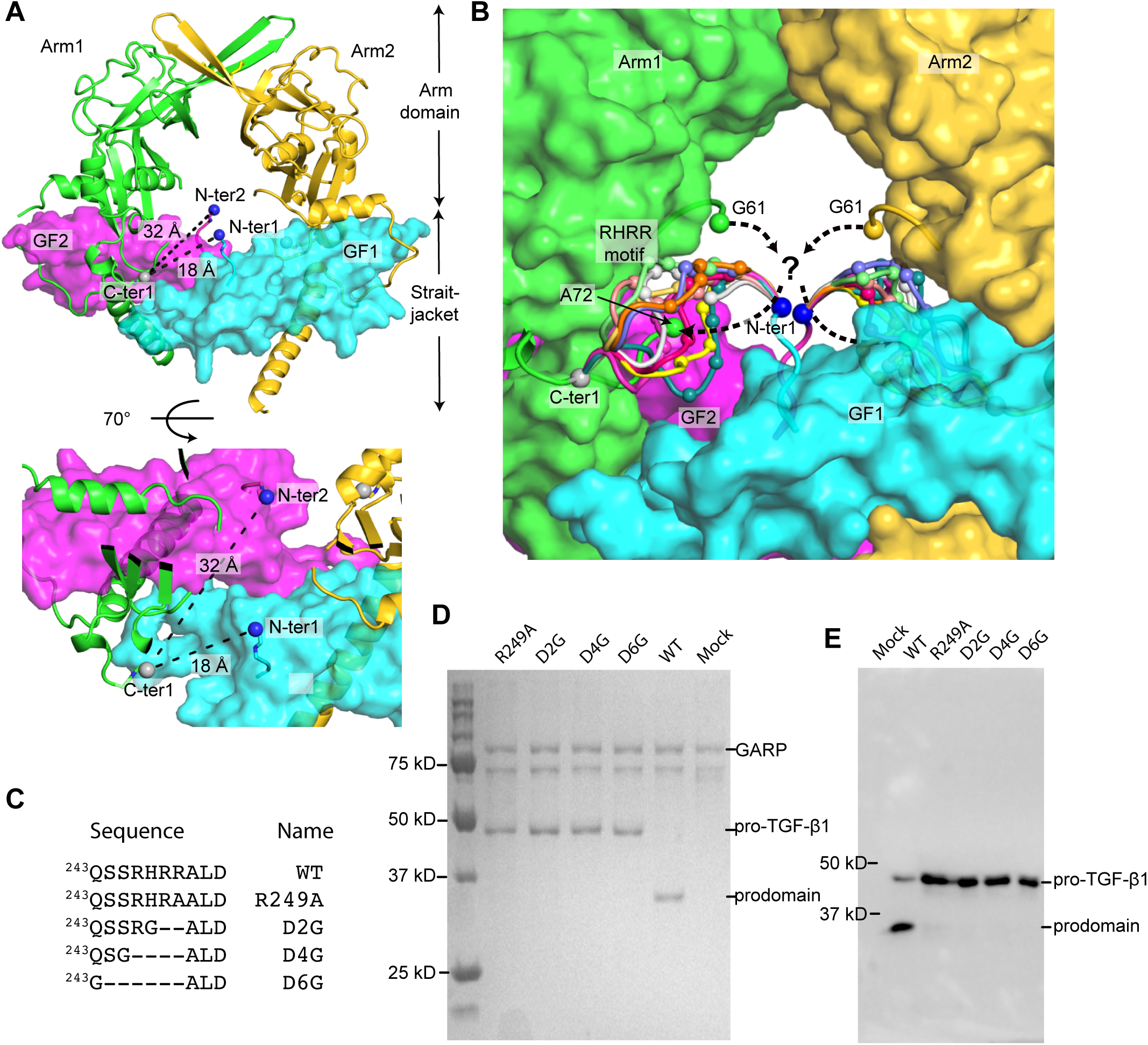
Prodomain-GF connections. **A.** Views of the two possible arm domain-GF connections, from residue 242 in one arm domain monomer (C-ter 1) to residue 251 in one GF monomer (N-ter 1) or the other monomer (N-ter 2). The gaps to be spanned are dashed. Spheres mark the C atom of C-ter 1 and N atoms of N-ter 1 and N-ter 2. Prodomain and GF domains are shown in cartoon and surface, respectively. **B.** Ten representative arm domain-GF connections (residues 243-250) built by MODELLER between the C-ter 1-N-ter 1 connection are shown as loops of different colors with Cαs of the RHRR PC cleavage site shown with small spheres. Residues 61 and 72, adjacent to the residues missing in density between the straitjacket and arm domain, are also marked with spheres (see Discussion). Remaining portions of pro-TGF-β1 are shown as surface. **C.** PC site mutations. **D.** WT and mutant pro-TGF-β1 co-expression with soluble GARP. Culture supernatants were incubated with StrepTactin Sepharose (GE healthcare) for 1 hour. StrepTactin beads were washed, heated at 100° C in reducing SDS sample buffer, and eluates subjected to SDS-PAGE and stained with Coomassie Brilliant Blue. **E.** Transient expression of WT and mutant pro-TGF-β1. Culture supernatants were subjected to reducing SDS-PAGE and Western blotting with antibody to the TGF-β1 prodomain (BAF246, R&D systems).

To further characterize the prodomain-GF connection, we shortened it by deleting 2 to 6 residues. We hypothesized that the physiologic connection should be flexible to enable PC cleavage. Thus, we expected that the N-ter 1-C-ter 1 connection should be capable of being shortened, without disrupting proper folding of pro-TGF-β1. Since mammalian cells have ER quality control systems that require proper protein folding prior to secretion, we assayed for the effect of deletion mutations on pro-TGF-β1 secretion by 293T cell transfectants. We deleted 2, 4, or 6 residues including the RHRR cleavage site, and introduced a glycine substitution to enable flexibility at the deletion position in the D2, D4, and D6 mutations (Fig. 3C). These mutations abolished PC cleavage and yet had no effect on level of pro-TGF-β1 expression (Fig. 3D, E). The ability to substantially shorten the linker without affecting expression, i.e. folding, supports the shorter of the two connections.

## Discussion

We describe here the crystal structure of a pro-TGF-β1 R249A mutant that is uncleaved by PC and contrast it with a re-refined structure of cleaved pro-TGF-β1. Clear differences are present adjacent to the cleavage site in at least 3 N-terminal GF residues. PC cleavage appears to result in no overall conformational changes in procomplex structure, although interesting differences are present distal from the cleavage site that appear to be influenced by lattice contacts that differ among structures. The R249A mutant is a model for the structure of the proform of pro-TGF-β1 present during biosynthesis, after folding is completed in the endoplasmic reticulum, and prior to cleavage by a PC protease in the Golgi. As such, it provides the closest glimpse yet available for the connectivities between prodomains and GF domains within individual pro-TGF-β1 monomers during biosynthesis.

SDS-PAGE of purified pro-TGF-β1 R249A mutant protein and crystals formed from it, along with the distinct conformation of the GF N-terminus, show that the prodomain-GF linker region was intact in this structure, although the electron density was too weak to be traced. Distance measurements and modeling based on this structure unambiguously define the shortest of two possible prodomain-GF connections as that which is physiologic. These results were further supported by the demonstration that shortening of the prodomain-GF linker by deletion of 2 to 6 residues had no adverse impact on the amount of pro-TGF-β1 biosynthesis or secretion. We therefore concluded that the prodomain arm domain is connected to the GF with which it has no noncovalent interaction (Fig. 4A). In contrast, the arm domain noncovalently interacts with the GF domain of the other monomer and forms a super β-sheet with it; the β1-strand of the arm domain jelly roll fold hydrogen bonds to the β7-strand of the β-finger of the GF domain from the other monomer (Fig. 2A).

**Figure 4:**
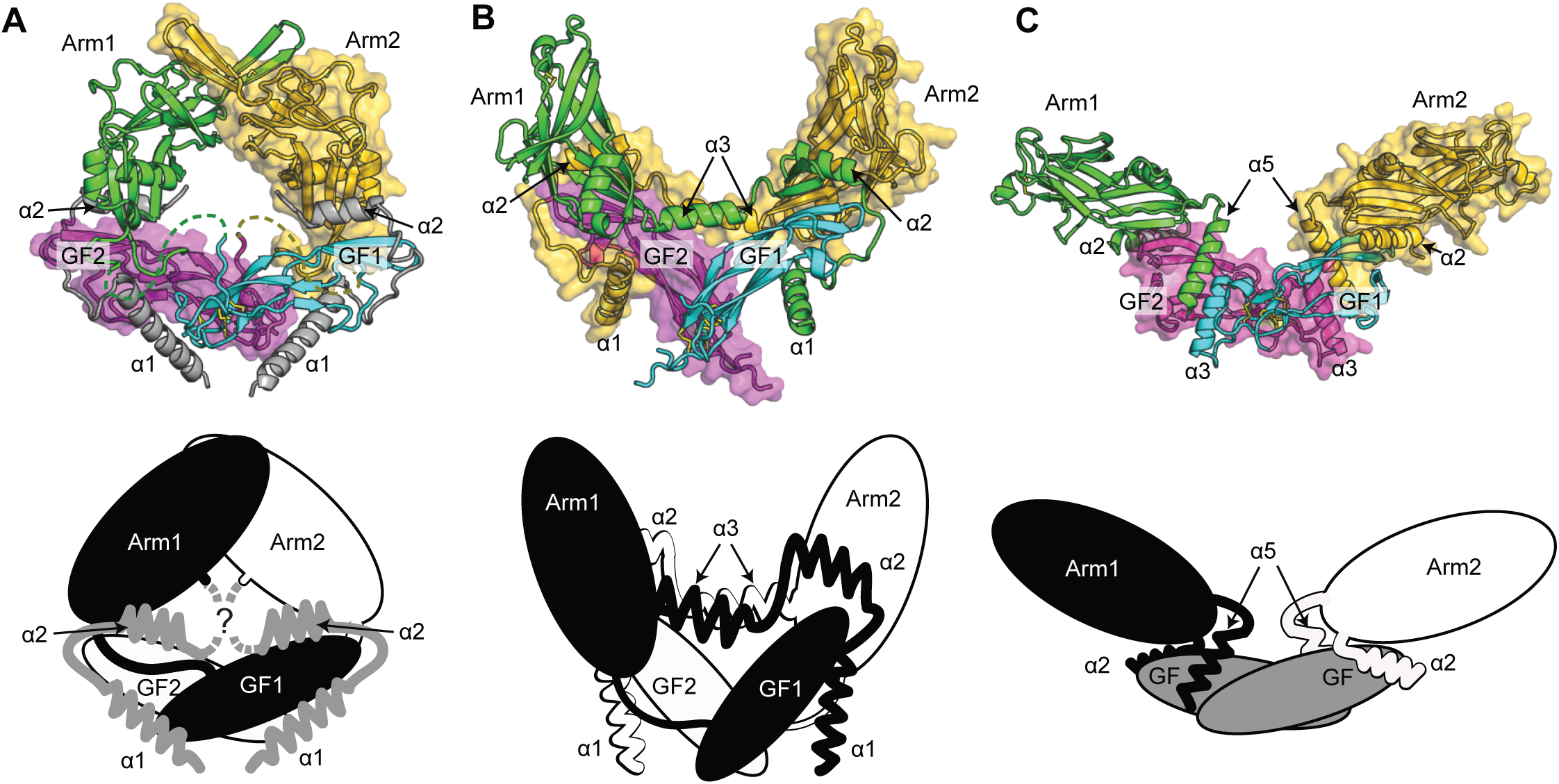
Prodomain-GF swapping. **A-C.** Prodomain-GF interactions of pro-TGF-β1 (**A**), pro-activin A (**B**) and pro-BMP9 (**C**). Upper: One putative monomer is shown in cartoon and the other as a surface. Lower: Schematic with one putative monomer in black and the other in white. The straitjacket region in pro-TGF-β1 (**A**) and the GF in pro-BMP9 (**C**) are in gray because the monomer to which they belong is unclear.

Comparison of our structures in regions distal from the PC cleavage site also provide insights into regions of the pro-TGF-β1 structure that are influenced by crystal lattice contacts and relevant to the flexibility of the prodomain in latency and activation. The jelly roll fold of the arm domain is invariant in structure, demonstrating it can provide a rigid pathway for transmission of force between the integrin bound to the RGDLATI motif in the shoulder of the arm domain to the opposite end of the arm domain that connects to straitjacket elements. The bowtie tail alters markedly upon integrin binding [19]. The association region covalently and noncovalently associates with milieu molecules that differ in 3D folds. Moreover, the 2:1 stoichiometry of pro-TGF-β monomer: milieu molecule complexes implies that the structure of the association region in each pro-TGF-β monomer must differ in complexes with milieu molecules [2]. In the lattices that enable crystallization of free pro-TGF-β1 to date, the bowtie tail and association region are prominent in lattice contacts, and have markedly different structures in an integrin-bound structure [19]. Furthermore, fewer residues in the association region are α-helical in the R249A mutant here than in the cleaved pro-TGF-β1 structure. The latency lasso also differs in conformation in the R249A mutant compared to the other structures as it wraps around the GF finger. This emphasizes the looseness of this straitjacket element. In contrast, structural comparisons show that the portion of the α1-helix that buries between the two GF monomers and the fastener that connects the arm domain to the α1-helix are highly conserved in position. This observation correlates with the finding that in molecular dynamic simulations of force-dependent integrin activation, this portion of the α1-helix and the fastener are more force-resistant than the latency lasso and association region [19].

It is interesting to examine our conclusions on prodomain-GF connectivity in pro-TGF-β1 in light of recent structures of pro-complexes for BMP-9 [23] and activin [24]. In pro-activin A, the PC cleavage site was replaced with a 3C protease site and a segment in the arm domain was deleted. Protein was refolded from E. coli, and with and without 3C cleavage, was subjected to crystallography. The prodomain-GF linker was disordered in both structures; however, based on the distance to be spanned, it was concluded that the topology was the same as described here. In other words, in both pro-TGF-β1 and pro-activin A, the arm domain connects to the GF with which it has no noncovalent interaction and intimately noncovalently interacts with the GF in the other monomer. In pro-TGF-β1 structures, the straitjacket shows density from the α1-helix through the latency lasso through to the end of the α2-helix. There is then a gap, with density missing in residues 62-71 that connect the straitjacket to the arm domain. The disordered residues lie in the central solvent-filled cavity in the pro-TGF-β1 ring (Fig. 3B and Fig. 4A lower). In pro-TGF-β1 crystal structures the straitjacket and arm domain that are closest to one another have been arbitrarily assigned the same chain IDs [14, 19]. However, the connection to the opposite arm domain on the other side of the cavity is also possible, since the distance to be spanned with or without crossing the channel is similar (Fig. 3B and 4A). Interestingly, in pro-activin A, the analogous region is a highly ordered helix (α3) and reaches across to the opposite arm domain so that the arm domain interacts with the α2-helix from the other monomer (Fig. 4B). Thus, there are two types of swaps in pro-activin A. In swap 1, the straitjacket elements including the α2-helix interact most closely with the GF from the same monomer and with the arm domain from the opposite monomer. In swap 2, the arm domain interacts most closely with the GF from the opposite monomer. We see swap 2 in pro-TGF-β1, and swap 1 is also possible. Interestingly, residues 62-71 missing in density in pro-TGF-β1 correspond in pro-TGF-βs 2 and 3 to a region with five additional residues, including one cysteine [1]. These cysteines should form an additional disulfide that dimerizes the prodomain. Perhaps structures of pro-TGF-βs 2 and 3 could resolve whether swapping occurs in this region.

Pro-BMP-9 [23] is yet another case. It lacks density for the α1-helix and latency lasso, yet shows good density for the α2-helix. Furthermore, the density between the α2-helix and the arm domain is continuous, and connects the α2-helix to the same arm domain with which it noncovalently associates (Fig. 4C). Thus, pro-BMP-9 lacks swap 1. Because the α1-helix and latency lasso are missing in density in pro-BMP9, could a putative swap 1 present in the pro-form have been reversed by conformational change after PC cleavage? The region responsible for swap 1 is in the loop between the straitjacket α2-helix and arm domain β1-strand. This loop is 12 residues longer in pro-TGF-β1 and 17 residues longer in pro-activin A than in pro-BMP-9 [2]. Even building a model of pro-BMP-9 with a conformation in which the two arm domains associate closely in pro-TGF-β1 [23], it appears that the loop between the α2-helix and β1-strand is not long enough for swap 1 to occur in pro-BMP-9. Thus in pro-BMP-9, swap 1 is neither observed experimentally nor feasible in an alternative pro-complex conformation. It thus appears that TGF-β family members differ in swapping among structural elements.

Swapping between the straitjacket, arm domain, and GF domain has profound biological implications for the TGF-β family at large. TGF-β family members form homodimers as well as heterodimers. Inhibin-β subunits homodimerize to form activins such as pro-activin A, as discussed above, and heterodimerize with inhibin-α subunits to form inhibins [12, 25, 26]. Some BMP heterodimers show higher activity than homodimers in vitro and in vivo, including BMP-4/7 [27-29] and BMP-2/7 [20, 30-32]. Although structures of biologically relevant heterodimers are not available, the structure here of uncleaved pro-TGF-β1 together with structures of pro-activin A and pro-BMP-9 suggest that swapping of two elements of the prodomain with the GF domain could provide mechanisms for preferential formation of heterodimers over homodimers for some TGF-β family members.

The TGF-β family is very diverse, with 33 genes giving rise to a larger number of homo and heterodimers that regulate all aspects of development and homeostasis. The TGF-β superfamily is larger than other extracellular protein families that regulate development including the Wnt, Frizzled, Delta/Jagged, and Notch families [2].

Multiple families that contain similar cystine-knot fold GF domains also emerged at the dawn of metazoans, but the TGF-β family multiplied more than any other. TGF-β has a larger prodomain than any other cystine-knot fold cytokine, and it has been argued that this size, combined with the capacity for diversification of the prodomain, contributed to the evolutionary success of the TGF-β family [2]. It has become increasingly clear that prodomains are of key importance in the physiology of the TGF-β family. The study here, and comparisons to other pro-complexes, extends our understanding of how TGF-β family prodomains interact with and regulate their GFs.

### Experimental procedures

**Protein expression and purification.** The human pro-TGF-β1 expression construct contains an N-terminal 8-His tag, followed by a SBP tag and a 3C protease site [14]. A C4S mutation and N-glycosylation site mutations N107Q and N147Q were introduced to facilitate protein expression, secretion and crystallization. In addition, R249A, D2G, D4G and D6G PC site mutations (Fig. 3C) were introduced. R249A mutant protein expression and purification were as described [19].

Human pro-TGF-β1 constructs with no N-terminal tags, WT Cys-4 and R249A, D2G, D4G, D6G PC site mutations (Fig. 3C) were also used for co-expression with GARP.

**Cell culture and transfection.** HEK293S GnTI^-^ stable cell lines expressing soluble GARP protein with N-terminal 6-His tag, SBP tag and a 3C protease site and HEK293T cells were maintained in DMEM supplemented with 10% fetal bovine serum (FBS), 4 mM L-glutamine, and 1% non-essential amino acids. Cells were transiently transfected with WT and mutant pro-TGF-β1 constructs using Lipofectamine 2000 (Invitrogen) according to the manufacturer’s instructions. Culture supernatants were harvested after 48 h to test for pro-TGF-β1 expression and co-expression with GARP.

**Crystal structures.** Crystals of pro-TGF-β1 R249A mutant (1 μl, 10 mg/ml, 20 mM Tris-HCl pH 8.0, 150 mM NaCl) were formed in hanging drops at 16 °C with 1 μl of 10% Dioxane, 100 mM MES pH 6.5, 1.7 M (NH_4_)_2_SO_4_ (well solution). Crystals were cryo-protected with well solution containing 31.4% Li_2_SO_4_. Diffraction data from GM/CA-CAT beamline 23-ID of the Advanced Photon Source (APS) at the Argonne National Laboratory were processed with XDS [33] with cross-correlation to determine the diffraction limit [34]. Structures were solved with molecular replacement by PHASER [35] with cleaved pro-TGF-β1 (PDB 3RJR) as the search model. The previous cleaved pro-TGF-β1 dataset [14] was reprocessed using XDS [33] with cross-correlation to determine the diffraction limit [34] to a resolution of 2.9 Å. Structures were refined with PHENIX [36], manually built with Coot, and validated with MolProbity [22].

**Modeling.** We used the MODELLER 9.12 loop modeling protocol [21] to build and refine the 8 missing residues between the prodomain and GF. Crystallographically defined regions were kept fixed. Ten loop models were built for each of the two connections, assessed based on the DOPE score [37], and validated using MolProbity [22].

